# Temporal changes in resting state networks induced by propofol anesthesia

**DOI:** 10.1101/2021.10.24.465655

**Authors:** Mi Kyung Choe, Seung-Hyun Jin, June Sic Kim, Chun Kee Chung

**Author notes:** **Competing Interest Statement:** The authors declare no competing interest.

## Abstract

The cerebral cortical changes associated with propofol-induced unconsciousness remain unknown. While the anesthetic agent affects the entire cerebral cortices, there might be spatiotemporal differences in cortical changes. In particular, we hypothesized that there might be spatiotemporal differences in cortical changes with propofol-anesthesia. To address this hypothesis, we investigated power spectrum changes in electrocorticography (ECoG) signals obtained during the induction phase from awake state to unconsciousness. We found that, 1) the power increased in the range of frequencies < 46 Hz (delta to low gamma), and decreased in the range (62–150) Hz (high gamma), in global channels during the induction phase. 2) The power in the frontoparietal network (FPN), specifically the superior parietal lobule and prefrontal cortex, started to change early, but took a long time to completely change. However, the power in the default mode network (DMN) started to change late, but took a short time to completely change. 3) The power change (*ΔPower*) in the DMN was more conspicuous than that of the dorsal attention network (DAN) in high gamma frequency. Considering that the FPN is involved in communication with the external world and that DMN is involved in communication with self, loss of consciousness induced by general anesthesia results from first, disrupted communication between self and external world, and is then followed by disrupted communication within self, with decreased activity of the FPN, and later, attenuated activity of the DMN.

**Significance Statement:** We investigated the spatiotemporal changes of power spectrum in human electrocorticography (ECoG) during the induction phase from awake state to unconsciousness. We found that from delta to low gamma frequency, the power increased, while in high gamma frequency, the power decreased over all channels. The power in the frontoparietal network (FPN) preferentially changed, then the power in the DMN changed later. The power in DMN decreased more than those in other RSNs in high gamma frequency. Loss of consciousness induced by general anesthesia results from first, disrupted communication between self and external world, followed by disrupted communication within self, with decreased activity of the FPN, and later, attenuated activity of the DMN.

## Introduction

Consciousness is one of the most daunting issues in neuroscience. Characterization of the changes in cortical activity that arise upon loss of consciousness induced by general anesthesia is important to understand the underlying mechanism of consciousness. General anesthesia is a reversible drug-induced condition that medically attains unconsciousness, amnesia, analgesia, and akinesia. Propofol is a commonly used anesthetic drug, and is known to induce an anesthetic state through enhancing inhibition at GABAA receptors (1). The molecular and cellular effects of anesthetics have been established, but their effects on the cortical activity remain unclear.

Anesthetic drugs cause global central nervous system suppression (2). However, anesthetic loss of behavioral response is not necessarily accompanied by an even block of all cortical regional activities. For example, during anesthesia, cortical response in the primary sensory cortex to sensory stimuli is preserved (3), while activities in the higher order cortical sensory area are suppressed (4). In previous studies investigating changes in cortical activities during unconsciousness by general anesthesia, it is unclear which regions are functionally related to loss of consciousness induced by general anesthesia.

Since most previous studies investigated cortical changes during steady state of general anesthesia, those approaches could not depict neurophysiological features at the onset of loss of consciousness from awake state. To specifically pinpoint crucial regions involved in consciousness, the identification of temporal and spatial changes in the activities of the human cortex are critical with tracking the transition into unconsciousness. We investigated the spatiotemporal dynamics of the cortex during the induction phase of propofol-induced anesthesia in human.

fMRI (functional MRI) and EEG (electroencephalography) are often used modalities to investigate the cortical activities during general anesthesia. However, the limitations of both modalities, i.e., fMRI in temporal resolution and EEG in spatial resolution, impeded unraveling the details of the transition into unconsciousness. Temporal information of the cortical activities from the awake state into unconsciousness is essential to unravel the details of transition. Electrocorticography (ECoG) is an invasive modality that can directly measure the neuronal population activities from the cortex. Hence, it has better temporal and spatial resolution than fMRI and EEG, respectively. Moreover, ECoG with high spectral resolution is compared to the EEG and fMRI, especially at high gamma frequency (> 50 Hz). It provides detailed information of human brain function with relatively high signal-to-noise ratio (SNR), and high sensitivity and temporal resolution (5, 6).

In this study, we investigated the temporal and spatial changes in the cortical areas during the induction into unconsciousness (hitherto, the ‘induction phase’). Patients with medically intractable epilepsy provide a unique opportunity to investigate ECoG during general anesthesia, since they need localization of the epileptogenic zone before resection surgery. We recorded ECoG in resection surgery to examine the spatiotemporal dynamics associated with loss of consciousness. Using power spectral analysis, we first investigated the spectral characteristics in the awake and unconscious state with propofol-induced general anesthesia.

Furthermore, we investigated spatiotemporal changes in the cortical areas during the induction phase of unconsciousness. For temporal changes, we assessed 1) the start point, and 2) the normalized time interval between the start and finish of power change (*Δnormalized t*). We also assessed 3) the power change (*ΔPower*) in the cortical areas. We hypothesized that during the induction phase of propofol anesthesia, the power in the cortical areas associated with consciousness would drastically change.

## Results

### Power difference between the awake state and unconscious state induced anesthesia

The power across range of frequencies < 46Hz increased in the unconscious state, while the power in the range (62–150) Hz decreased in the unconscious state (Fig. 1).

**Figure 1.**
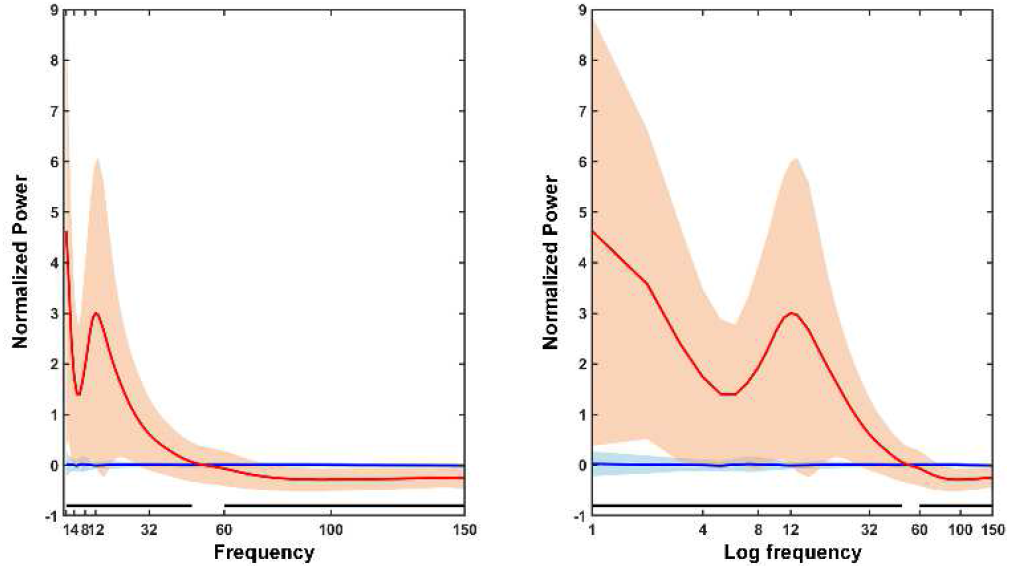
Electrocorticographic (ECoG) power spectrum over increasing frequency frequencies. Red line indicates the average of normalized power during the unconscious state. The blue line indicates the average of normalized power while awake. The shaded area indicates standard deviation. Black line indicates statistically significant power difference per frequency bin (p < 0.01/82).

### The power difference of the cerebral cortex and resting state networks

The power in all cortical regions except the secondary visual cortex (V2) increased in the range < 46 Hz. In the range (62–150) Hz, the power in all cortical regions decreased (Fig. S1 of the Supplementary Information (SI)). Moreover, the power in the range of frequencies < 46 Hz increased in all RSNs. The power in all RSNs decreased in the range (62–150) Hz (Fig. 2).

**Figure 2.**
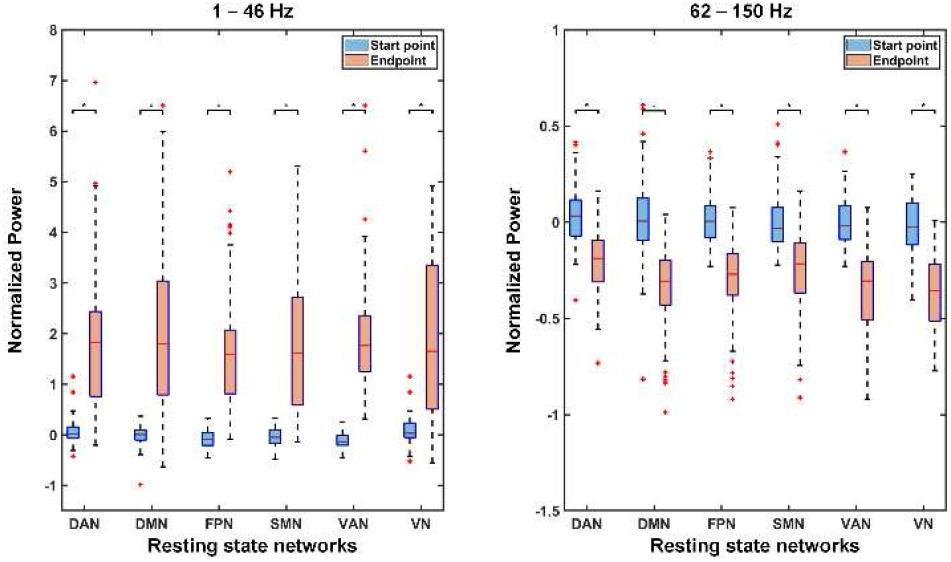
The normalized power of resting state networks at the start point and endpoint. Upper: Range of frequencies < 46Hz; Lower: Range (62–150) Hz. * p < 0.05/6. Abbreviations: DAN = Dorsal attention network; DMN = Default mode network; FPN = Frontoparietal network; SMN = Sensorimotor network; VAN = Ventral attention network; VN = Visual network.

### The start point of power change across the cortex

In the range of frequencies < 46 Hz, the normalized time point of the start of power change in SPL (part of the DAN and the FPN) was earlier than those in global channels and the DMN (angular gyrus, AnG, the opercular and triangular part of the inferior frontal gyrus, OpTr, middle temporal gyrus, MTG, superior temporal gyrus, STG, and temporopolar area, TPC). The start point of power change in SPL was earlier than those in the somatosensory cortex (S1), part of the SMN, and associative visual cortex (V3–5), part of the VN. The start point of power change in SPL was also earlier than those in the frontal eye fields (FEF), and dorsolateral prefrontal cortex (DLPFC). The start point of power change in STG and AnG, part of the DMN, were later than those in the global channels, DLPFC and the supramarginal gyrus (SMG), part of the FPN. The start point of power change in V3–5, part of the VN, was later than those in the global channels, DLPFC and SMG, part of the FPN. The start point of power change in STG was later than that in the anterior prefrontal cortex (aPFC) (Table S1 & Fig. S2 of the SI). In the range (62–150) Hz, the start point of power change in SPL, part of the FPN was earlier than those in global channels, the VN (V3–5), and the DMN (MTG, STG, and AnG). The start point of power change in AnG, part of the DMN, was later those in the global channels, DLPFC and SMG, part of the FPN. The start point of power change in AnG was also later than those in aPFC and STG (Table S1 & Fig. S2 of the SI).

### The start point of power change within the resting state networks

In the range of frequencies < 46 Hz, the start point of power change in the DMN was later than those in the FPN, VAN, and SMN. The start point of power change in the VN was later than those in the DAN, and FPN. In the range (62–150) Hz, the start point of power change in the FPN was earlier than that in the DMN, SMN, and VN. (Table 1, Fig. S3 of the SI).

**Table 1.** Statistical significance of the start point of power change within the resting state networks at the induction phase of unconsciousness.

| Frequency | RSN 1 | RSN 2 | Mean difference | P |
| --- | --- | --- | --- | --- |
| < 46Hz | DMN | DAN | DMN > DAN | 0.00002 |
|  |  | FPN | DMN > FPN | 0.00000 |
|  |  | SMN | DMN > SMN | 0.00284 |
|  | VN | DAN | VN > DAN | 0.00248 |
|  |  | FPN | VN > FPN | 0.00002 |
| > 62Hz and < 150Hz | FPN | DMN | FPN < DMN | 0.00000 |
|  |  | SMN | FPN < SMN | 0.00250 |
|  |  | VN | FPN < VN | 0.00044 |

**Abbreviations:** DAN = Dorsal attention network; DMN = Default mode network; FPN = Frontoparietal network; SMN = Somatomotor network; VN = Ventral attention network.

### The difference of the *Δnormalized t* across the cortex

In the range of frequencies < 46 Hz, the *Δnormalized t* in SPL (part of the FPN and DAN) was longer than those in the global channels, and the DMN (MTG, STG, TPC, AnG, and OpTr). The *Δnormalized t* in SPL was also longer than those in V3–5, part of the VN, and FEF (Table S2 & Fig. S4 of the SI). In the range (62–150) Hz, the *Δnormalized t* in DLPFC, part of the FPN was longer than those in the global channels, the DMN (MTG, STG, and AnG), SMN (S1), and the VN (V2 and V3–5). The *Δnormalized t* in DLPFC was also longer than those in SPL and SMG. The *Δnormalized t* in AnG, part of the DMN, was shorter than those in the global channels, supplementary motor area and premotor cortex (SMA), and STG. The *Δnormalized t* in aPFC was longer than those in the global channels, S1, SPL, V3–5, MTG, STG, and AnG. The *Δnormalized t* in S1 was shorter than those in the global channels (Table S2 & Fig. S4 of the SI).

### The difference of the *Δnormalized t* within the resting sate networks

In the range of frequencies < 46 Hz, the *Δnormalized t* in the FPN was longer than those in the DMN and VN. The *Δnormalized t* in the DMN was shorter than that in the DAN. In the range (62–150) Hz, the *Δnormalized t* in the FPN was longer than those in the DAN, DMN, SMN, and VN. The *Δnormalized t* in the DMN was longer than that in the VN (Table 2, Fig. S5 of the SI).

**Table 2.**
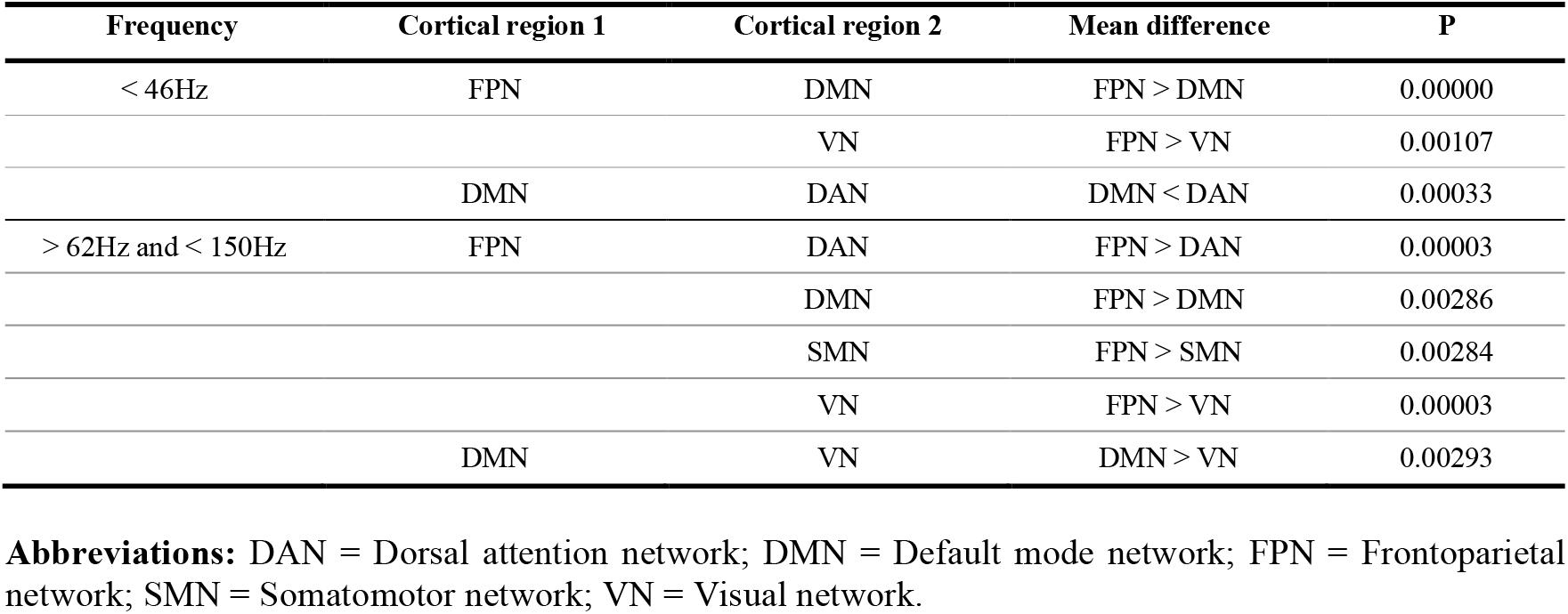
Statistical significance of the *Δnormalized t* within the resting state networks at the induction phase of unconsciousness.

### The difference of *ΔPower* across the cortex

The power across the range of frequencies < 46 Hz in V2 increased less than those in the global channels and OpTr. In the range (62–150) Hz, the power in S1 decreased less than those in the global channels, AnG, and DLPFC. The power in SPL involved in the FPN and DAN decreased less than those in the global channels, the DMN (aPFC, STG, AnG, and OpTr), V2, V3– 5, and DLPFC. The power in AnG decreased more than the power in MTG (Table S3 & Fig. S6 of the SI).

### The difference of the *ΔPower* within the resting state networks

There was no significant difference of the *ΔPower* within RSNs across the range of frequencies < 46Hz. The power in the range (62–150) Hz in the DMN decreased more than the power in the DAN. The power in the VAN decreased more than the power in the DAN (Table 3, Fig. S7 of the SI).

**Table 3.**
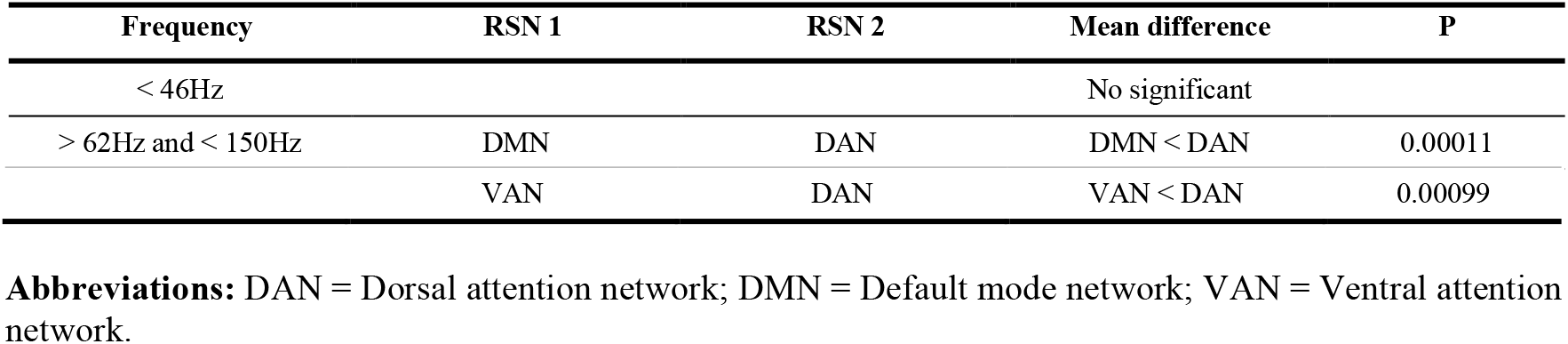
Statistical significance of the *ΔPower* within the resting state networks at the induction phase of unconsciousness.

## Discussion

In the present study, we investigated the temporal and spatial characteristics of power changes in the cortical regions with the relevant resting state networks tracking the transition into unconsciousness. The power change was not uniform across diverse frequencies and cortices. The power increased from delta to low gamma frequency, while the power decreased in high gamma frequency in global channels. The power significantly increased from delta to low gamma in all cortical regions except V2 and in all RSNs, whereas it decreased in high gamma in all cortical regions and all RSNs. Second, the temporal changes between FPN, DMN, and VN are different during the induction phase. In the start point and Δnormalized t, the power in FPN, specifically the superior parietal lobule (SPL; BA7) and prefrontal cortex (DLPFC; BA9), started to change early, but took a long time to complete the change. However, the power in DMN and VN started to change late, but took a short time to complete the change. Third, the ΔPower in DMN and VAN was more conspicuous than that of DAN in high gamma frequency.

In the present study, we identified the increased power across the range of delta to low gamma, and decreased power in high gamma frequency. Previous EEG studies indicated that with increasing propofol concentration, the power spectrum shifts from a high-frequency, low-amplitude activity to a low-frequency, high-amplitude activity (5, 7). However, power changes in beta of (13–30) Hz and low gamma of (25–40) Hz frequency during propofol induction were inconsistent with previous studies (5, 8-10). In the present study, we found that in the transition from awake state to unconscious state, power in the global ECoG channels increased at lower frequency (< 46 Hz), and decreased at high gamma frequency. The decreased high gamma power in the local field potential (LFP) was accompanied by decreased action potential firing rate in the neocortex, with increasing propofol concentration (7). The shift towards lower frequency indicates thalamic hyperpolarization. The coupling between gamma power and the phase of delta frequency was enhanced by the induced propofol anesthesia, indicating the hyperpolarization of the cortical and subcortical circuit (5, 11). Taken together, GABAA inhibition induced by propofol alters the integration of the excitatory and inhibitory inputs of neurons, decreasing action potential firing and the balance of excitatory and inhibitory postsynaptic potentials in the cortical and subcortical circuits, increasing power at lower frequency and decreasing power at high gamma frequency.

Consciousness has two key components: 1) wakefulness (i.e., the level of consciousness), and 2) awareness (i.e., the content of consciousness) (12). Awareness consists of external and internal awareness. External awareness indicates the perception of environmental stimuli such as seeing, tasting, smelling, and hearing. Internal awareness means external stimuli-independent thoughts such as emotion, self-reference, and inner speech (13, 14). Staying consciousness refers to being able to maintain wakefulness, awareness of the external world, and awareness of self (15). In other words, loss of consciousness would result from communication loss between self and external world, and within self.

Figure 3 is a graphical representation of the temporal change of power at lower frequency (< 46 Hz) and at high gamma frequency of (62–150) Hz during the induction phase of unconscious in RSNs and the cortex (Fig. 3). It summarizes the main findings of the present study, indicating how fast (the start point and Δnormalized t) and by how much (ΔPower) the power within RSNs changed in lower and high gamma frequency.

**Figure 3.**
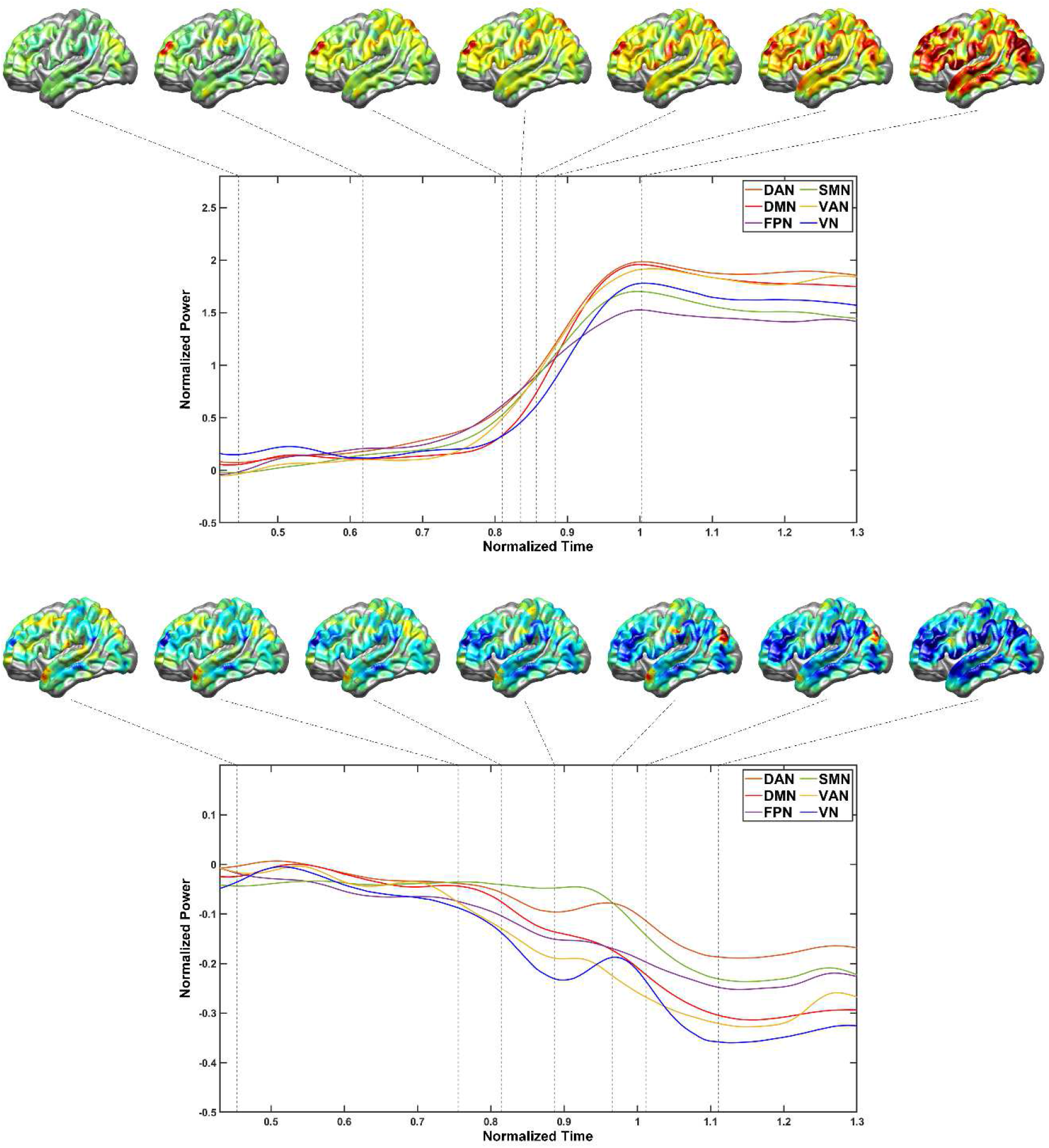
Temporal change during the induction phase of unconsciousness in RSNs and the cortex. Upper: The range of frequencies < 46 Hz; Lower: The range (62–150) Hz. Abbreviations: DAN = Dorsal attention network; DMN = Default mode network; FPN = Frontoparietal network; SMN = Sensorimotor network; VAN = Ventral attention network; VN = Visual network.

The present study highlights the temporal characteristics of power change within RSNs and the cortical regions, which have not been observed in previous studies. The FPN started to change early, but took more time to completely change, whereas the DMN started to change late, but took less time to completely change. That is, the propofol anesthesia preferentially affects the activities of the FPN earlier, then affects the activities in the DMN later. The FPN is engaged in the external awareness and perception of information from the environment (16). Specifically, the superior parietal lobule (SPL, BA7), dorsolateral prefrontal cortex (DLPFC, BA9, 46), and inferior parietal lobule (SMG, BA40) that started to change earlier in the present study are involved in conscious awareness (17, 18), including attention, sensory perception, and integrating inputs from unimodal sensory regions (19-22). The DMN is the crucial network of self-awareness and internal awareness (23). Therefore, the external awareness would be disrupted with the decreased activity of the FPN during the loss of consciousness induced by general anesthesia. With the foremost external awareness disrupted, the internal awareness would begin to be disrupted with the attenuation of the DMN activity (Fig. 3).

The present study identified the difference of power change (ΔPower) within RSNs at lower frequency (< 46 Hz), and high gamma frequency of (62–150) Hz. The power in DMN (aPFC, STG, AnG, and OpTr) decreased more than those in other RSNs at high gamma frequency. The DMN is the dominant network associated with spontaneous cognition, and is most active during the time when people do not engage in external tasks (24, 25). The DMN engages in self-related mental activity, integrating information processing across the brain (25). The relatively reduced activity in the DMN, thought to be dominant in passive resting state condition, indicates the collapsed awareness of self (Fig. 3).

In high gamma frequency, the power in S1 was also relatively less decreased. In previous studies, the activities in low-level sensory cortices were relatively preserved (3, 16). However, the power across the range of frequencies < 46 Hz in V2 was relatively less changed, whereas more decreased at high gamma frequency. A previous study indicated that the power in high gamma conveyed maximal visual processing (26). Therefore, the power of high gamma in the visual cortex may be more decreased than those in other regions.

ECoG provides high quality neural population signals from the cortex directly, with better temporal resolution compared to non-invasive measurements (5, 27). In the present study, the mean time to loss of consciousness was 195.1 s (see Table S4 of the SI). During the first few minutes after starting the infusion of propofol, neural activities in all RSNs were changed. The high temporal resolution of ECoG showed how the power in RSNs correlated with loss of consciousness changed within a few seconds. The SNR for high gamma frequency of ECoG is greater than that of EEG, because the skull acts as a low-pass filter, and artifacts from the cranial muscle affect the SNR for high gamma frequency (28). In the present study, we identified the activity changes of RSNs in high gamma frequency, for which non-invasive measurements were hard to detect.

There are several limitations that need to be considered in interpreting the present results, and that should be addressed in future research. The spatial distributions of electrodes were uneven for each subject, since the electrode coverage was determined solely by clinical considerations. We could not record the ECoG from all cortical areas, nor evenly in the cortical regions. Therefore, the results of the present study can be interpreted in the context of the only networks and cortical regions covered in this study. Future studies should warrant more details for spatial difference with cortical distribution.

## Materials and Methods

### Participants

From 70 patients with medically intractable epilepsy who agreed to participate, we selected 27 patients with ECoG recording analyzable for the induction of general anesthesia using propofol. Excluded were 43 patients without analyzable ECoG during the induction phase. Also, we excluded 9 patients with abundant artifacts and 2 without behavior data, and finally, we analyzed in this study the ECoG data of 16 patients (8 males and 8 females; age: mean = 33.1, SD = 12.3 years; 8 right and 8 left hemispheres, Table S4 of the SI for details, and Fig. S8 of the SI). Patients were implanted with subdural grids and strips or depth electrodes for sole clinical purposes. One patient was implanted on two separate occasions for clinical necessity. This study protocol was approved by the Institutional Review Board at Seoul National University Hospital (H-1405-025-577). All participants in the study provided written informed consent.

### Anesthesia

The recordings were obtained during the induction phase of propofol-induced anesthesia for resection surgery after localization of the epileptogenic zone with ECoG electrodes implanted. Before propofol infusion, the recordings in the awake state were obtained (Average recording time during awake state, 204.2 s; standard deviation, 166.4 s). Thereafter, anesthesia was induced using target-controlled infusions of propofol, which continued until subjects lost the ability to respond to verbal commands (Average recording time from starting infusion to loss of verbal responsiveness, 195.1 s; standard deviation, 61.8 s).

### Electrode Localization

Patients had been implanted with subdural ECoG, depth electrodes (Ad-Tech Medical, Racine, WI, USA and PMT, Chanhassen, MN, USA) or high-density ECoG (PMT, Chanhassen, MN, USA). Subdural electrodes were of 4 mm diameter with 10 mm inter-electrode distance, while high-density ECoG were of 2 mm diameter with 4 mm inter-electrode distance. Depth electrodes had a surface area of 0.059 cm2 with 6 mm inter-electrode distance. Preoperative MR images were acquired using a Magnetom Trio Tim 3T scanner (Siemens, Erlangen, Germany) or Signa 1.5-T scanner (GE, Boston, MA, USA). Postoperative CT images were acquired using a Somatom sensation device (64 eco; Siemens München, Germany). Preoperative MR data were co-registered to postoperative CT images using CURRY software (versions 7.0 and 8.0; Compumedics Neuroscan) to localize the electrode locations of individual subjects.

The location of individual coordinates was converted to MNI coordinates and also projected onto the MNI surface template consisting of 81,924 nodes using CIVET pipeline (ver. 1.1.7, MNI) (29, 30). The location of the nearest node on the cortical surface was used based on the Euclidean distance for each electrode location. We also constructed the surface map of electrode coordinates to Brodmann areas defined in MNI coordinates (Fig. 4, Table S5 of the SI) (31, 32). We flipped the electrodes located in the right hemisphere to the corresponding nodes in the left hemisphere, because in this study, we focused on the cortical characteristic within the hemisphere, rather than the hemisphere difference. We grouped cortical areas into six resting state networks (RSNs) to compare the spatial distributions of RSNs (Table 4), including the dorsal attention network (DAN), the default mode network (DMN), the frontoparietal network (FPN), the somatomotor network (SMN), the ventral attention network (VAN), and the visual network (VN) (31-38). The cortical areas in RSNs are not mutually exclusive, since some cortical regions are parts of several RSNs. For example, SMG is part of VAN and FPN (33, 34, 36).

**Table 4.** Cortical regions of the resting state networks.

| Network | Cortical regions | Abbreviations | Number of Electrodes | Reference |
| --- | --- | --- | --- | --- |
| **DAN** | Supplementary motor area and premotor cortex | SMA | 91 | Aboitiz et al., 2014 (34) |
|  | Superior parietal lobule | SPL |  |  |
|  | Frontal eye fields | FEF |  |  |
|  | Associative visual cortex | V3–5 |  |  |
| **DMN** | Anterior prefrontal cortex | aPFC | 184 | Smallwood et al., 2021 (36)<br>Andrews-Hanna et al., 2014 (43) |
|  | Middle temporal gyrus | MTG |  |  |
|  | Superior temporal gyrus | STG |  |  |
|  | Temporopolar area | TPC |  |  |
|  | Angular gyrus | AnG |  |  |
|  | The opercular part and triangular part of the inferior frontal gyrus | OpTr |  |  |
| **FPN** | Superior parietal lobule | SPL | 139 | Nekovarova et al., 2014 (33)<br>Smallwood et al., 2021 (36) |
|  | Dorsolateral prefrontal cortex | DLPFC |  |  |
|  | Supramarginal gyrus | SMG |  |  |
| **SMN** | Primary somatosensory cortex | S1 | 62 | Chenji et al., 2016 (38) |
|  | Supplementary motor area and premotor cortex | SMA |  |  |
|  | Primary gustatory cortex | GC |  |  |
| **VAN** | Supramarginal gyrus | SMG | 53 | Aboitiz et al., 2014 (34) |
|  | The opercular part and triangular part of the inferior frontal gyrus | OpTr |  |  |
| **VN** | Secondary visual cortex | V2 | 38 | Yeo et al., 2011 (35) |
|  | Associative visual cortex | V3–5 |  |  |

**Abbreviations:** DAN = Dorsal attention network; DMN = Default mode network; FPN = Frontoparietal network; SMN = Somatomotor network; VAN = Ventral attention network; VN = Visual network.

**Figure 4.**
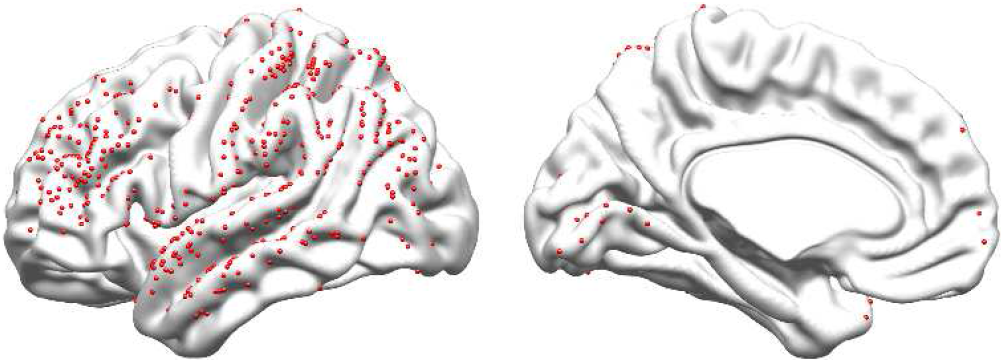
Locations of ECoG electrodes pooled in all patients analyzed. The locations of the 440 ECoG electrodes put on the left hemisphere for visual purpose are depicted by red circles on the brain template. The electrodes in the right hemisphere were flipped to the corresponding electrodes in the left hemisphere.

### Data Acquisition and Preprocessing

The recordings were obtained using SynAmps2 (Neuroscan) data acquisition system. All analyses were performed using MATLAB R2020a (MathWorks, Natick, MA, USA). ECoG data were sampled at 2,500 Hz, and down-sampled to 500 Hz. ECoG data were further frequency-pass filtered at (0.1 – 200) Hz. Channels disconnected from the amplifier or corrupted with line noise were excluded (39). The segment of 60 s artifact-free during the awake state was used for time–frequency analysis after artifacts rejection, except one with 20 s artifact-free segment. The signals were notch filtered at (60, 120, and 180) Hz, and were re-referenced to the common average reference (CAR).

### Time–Frequency Analysis

The complex Morlet wavelet transform was applied for time–frequency analysis. We defined the baseline period as (0 to 30) s in the awake state to calculate the normalized power. The absolute values of transformed data were normalized by the mean and standard deviation of the baseline period (Fig. 5). The power difference between unconsciousness and awake was calculated to determine the characteristic of power change in each frequency. The average power spectra during the awake period (60 s while awake) were compared with the average power spectra during the unconsciousness period (60 s after loss of response to verbal commands), by taking the average power over all channels in this study. We computed the mean difference between periods for each frequency using paired t-tests with Bonferroni correction (p < 0.01/82).

**Figure 5.**
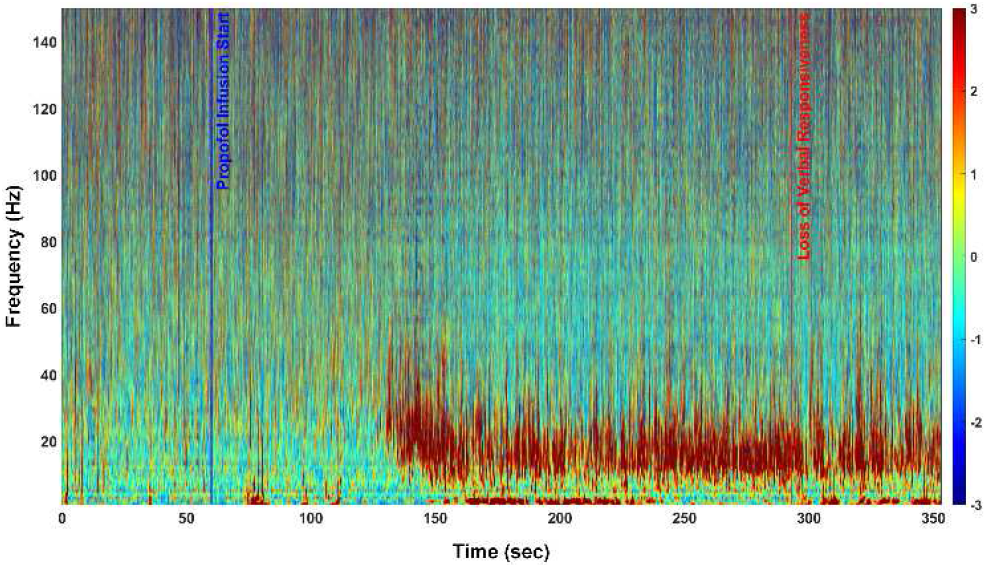
Representative time–frequency spectrogram during the induction of propofol anesthesia. Black line indicates the starting time of propofol infusion and the time of loss of verbal responsiveness. The color bar indicates the normalized power.

### Decreased peak frequency during the induction phase

To characterize the transition of the peak frequency during the induction phase (40), we calculated the average peak frequency in the awake and unconsciousness states. The peak frequency decreased from (23 to 12) Hz during the induction phase. We also estimated the spatial pattern of power at the peak frequency. We found that the frontal peak frequency shifted from beta (23 Hz) to alpha (12 Hz) frequency (Fig. S9 of the SI).

### Time Normalization

The time at the loss of verbal responsiveness during the administration of propofol was variable from subjects, due to individual variability (41). We normalized the temporal data from each subject to the same format. We defined the normalized time 0 as the time when propofol infusion started. We defined the normalized time 1 as the time when the change in average power spectrum of all channels in frequencies < 46 Hz finished, since low-frequency power increase was accompanied by the loss of behavioral response (40).

We defined the normalized time 1 based on power change, rather than verbal responsiveness, because we evaluated the patients’ response to verbal stimulation intermittently. Loss of response to verbal stimulation did not precede the power change in any subject. Normalized time of the data from the original range was rescaled using min–max normalization, so that all values were within the range 0 (tanes start) to 1 (tchange finish), as follows:

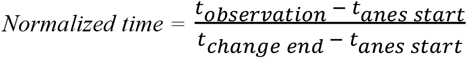

### Estimating temporal changes of power in the cortical areas

To effectively estimate the changing patterns of power during the loss of consciousness, smoothed power was calculated using Gaussian Filter with window length of 30 s (42). To investigate the critical regions of resting state networks changes accompanied by loss of consciousness, we estimated 1) the start point, 2) the normalized time interval between the start and finish of power change (Δnormalized t), and 3) the power change (ΔPower) in the cortical areas (Fig. 6). Kruskal–Wallis test and Mann–Whitney test were performed for statistical analyses. The significant difference was set at corrected p < 0.05/105.

**Figure 6.**
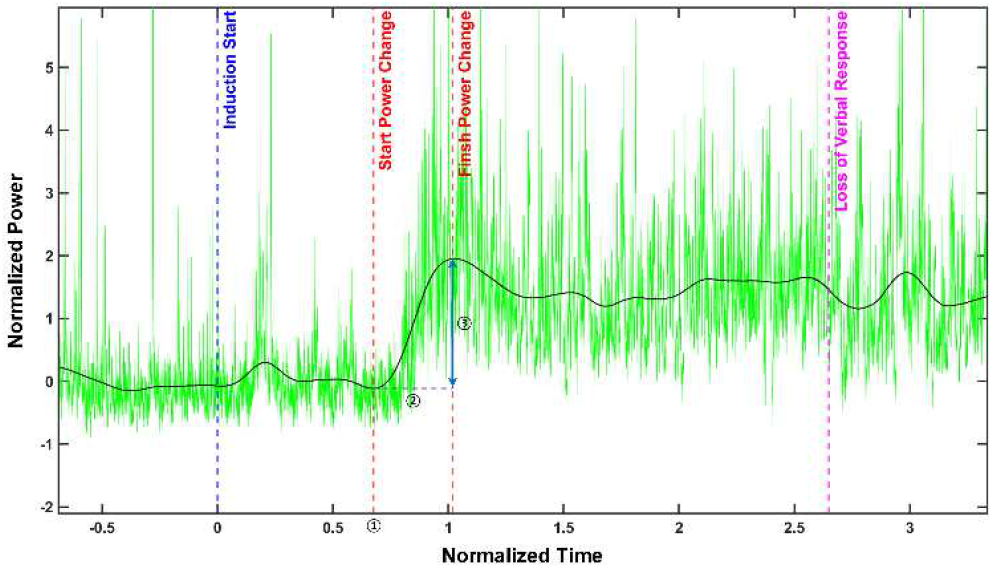
Representative example of estimating the start point, *Δnormalized t*, and *ΔPower* (< 46 Hz). Red vertical dashed line indicates the start point (➀) and the endpoint of power change. Purple horizontal dashed line indicates the normalized time from the start to finish of power change (*Δnormalized t*, ➁). Blue vertical line with two arrow heads indicates the power change (*ΔPower*, ➂). Magenta vertical dashed line indicates the time point of the loss of verbal responsiveness. Blue vertical dashed line indicates the time point of the start of propofol infusion.

#### The Start point

We defined the start point of power change as the normalized time point at which the gradient sign changes from negative to positive (< 46Hz), or changes from positive to negative in the range (62–150) Hz when the smoothed power starts to change.

#### The normalized time interval between the start and finish of power change (Δnormalized t)

To estimate the normalized time interval from the start point to the endpoint, we defined the endpoint of power change. We defined the endpoint of power change as the normalized time point at which the gradient sign changes from positive to negative (< 46 Hz), or changes from negative to positive in the range (62–150) Hz when the smoothed power finished changing. Furthermore, we calculated the normalized time interval between the start and finish of power change (Δnormalized t = the normalized time at endpoint – the normalized time at start point).

#### The Power change (ΔPower)

We calculated the power change (ΔPower) as the power difference between the power at the start point and endpoint (ΔPower = the normalized power at endpoint – the normalized power at start point).

### Estimating the temporal changes of power in the resting state network (RSNs)

We grouped cortical areas into six RSNs, not mutually exclusive, to compare the spatial distributions of RSNs changes. We estimated 1) the start point, 2) the Δnormalized t, and 3) the ΔPower in the RSNs (Fig. 6). Independent two-sample t tests were performed for statistical analyses. The significant difference was set at corrected p < 0.05/15.

## Acknowledgments

The authors thank Professor Hee-Pyoung Park of the anesthesiology department for help with data acquisition and providing valuable comments for study design. We are grateful to the intraoperative neuromonitoring technologists Gil Ho Kwak, Bo Eun Kim, Hyoung Jin Kim, Ji Hyang Nam, Jeongeum Park, and Young-Doo Choi for technical support in data acquisition.

This research was supported by the Basic Science Research Program through the National Research Foundation of Korea (NRF) funded by the Ministry of Education (2015R1D1A1A02061486) and the Ministry of Science & ICT (2019R1A2C1009674), South Korea.

## Author Contributions

S.-H.J., M.K.C., and C.K.C. designed the study and collected the data; M.K.C., J.S.K., and C.K.C. performed analyses and interpretation of data. M.K.C., J.S.K., and C.K.C. wrote the paper.

