## Supplementary Information for "Temporal changes in resting state networks induced by propofol anesthesia"

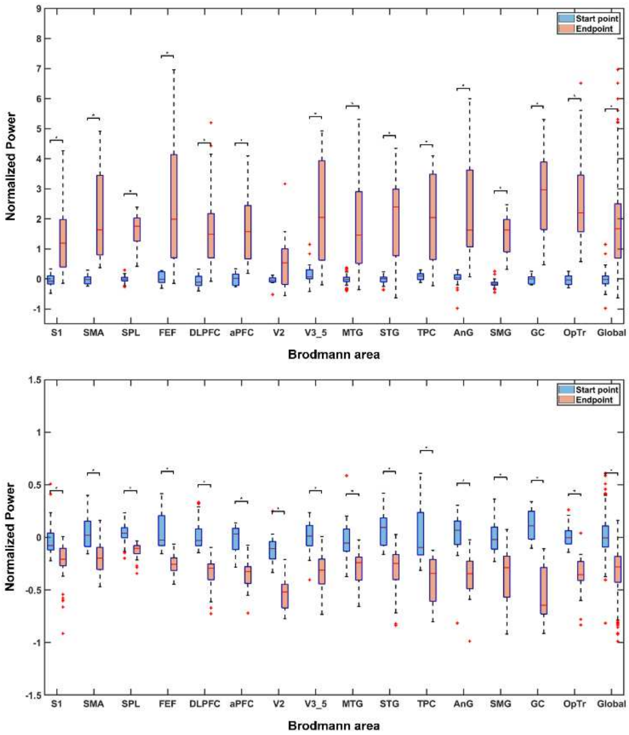


**Fig. S1. The normalized power across the cortex at start point and end point.** Upper: Range of frequencies < 46 Hz; Lower: Range of (62–150) Hz. * $p < 0.05/15$. Abbreviations: S1 = Primary somatosensory cortex (BA 1, 2, 3); SMA = Supplementary motor area and premotor cortex (BA 6); SPL = Superior parietal lobule (BA 7); FEF = Frontal eye fields (BA 8); DLPFC = Dorsolateral prefrontal cortex (BA 9, 46); aPFC = Anterior prefrontal cortex (BA 10); V2 = Secondary visual cortex (BA 18); V3–5 = Associative visual cortex (BA 19); MTG = Middle temporal gyrus (BA 21); STG = Superior temporal gyrus (BA 22); TPC = Temporopolar area (BA 38); AnG = Angular gyrus (BA39); SMG = Supramarginal gyrus (BA40); GC = Primary gustatory cortex (BA 43); OpTr = The opercular part and triangular part of the inferior frontal gyrus (BA 44, 45); Global = all ECoG channels.

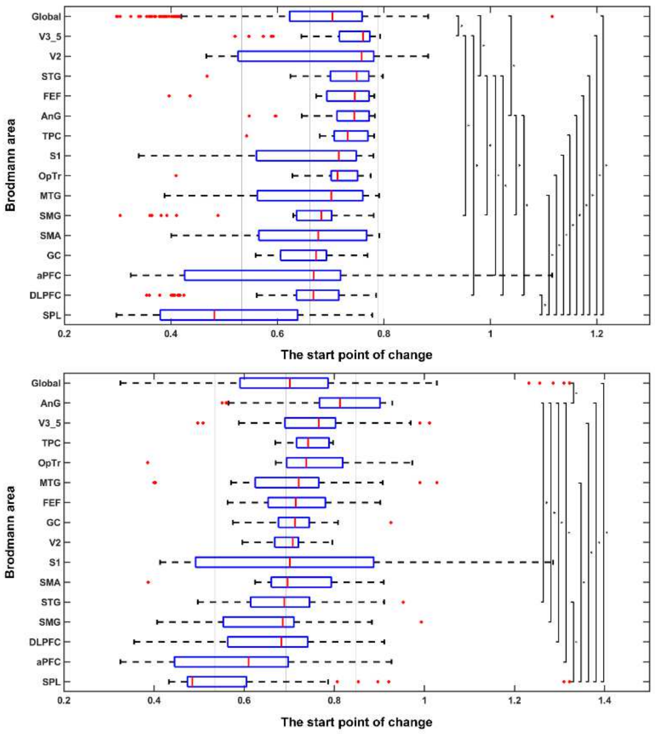


**Fig. S2. Boxplot of the start point of power change across the cortex at the induction phase of unconsciousness.** Upper: Range of frequencies < 46 Hz; Lower: Range of (62–150) Hz. * $p < 0.05/105$. Upper reference line = mean + standard deviation of all channels; Middle reference line = mean of all channels; Lower reference line = mean – standard deviation of all channels. Abbreviations: S1 = Primary somatosensory cortex (BA 1, 2, 3); SMA = Supplementary motor area and premotor cortex (BA 6); SPL = Superior parietal lobule (BA 7); FEF = Frontal eye fields (BA 8); DLPFC = Dorsolateral prefrontal cortex (BA 9, 46); aPFC = Anterior prefrontal cortex (BA 10); V2 = Secondary visual cortex (BA 18); V3–5 = Associative visual cortex (BA 19); MTG = Middle temporal gyrus (BA 21); STG = Superior temporal gyrus (BA 22); TPC = Temporopolar area (BA 38); AnG = Angular gyrus (BA39); SMG = Supramarginal gyrus (BA40); GC = Primary gustatory cortex (BA 43); OpTr = The opercular part and triangular part of the inferior frontal gyrus (BA 44, 45); Global = all ECoG channels.

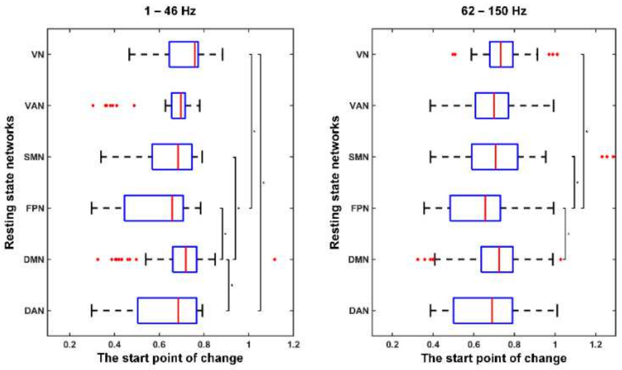


**Fig. S3. Boxplot of the start point of power change within resting state networks at the induction phase of unconsciousness.** Left: Range of frequencies < 46 Hz; Right: Range of (62–150) Hz. * p < 0.05/15. Abbreviations: DAN = Dorsal attention network; DMN = Default mode network; FPN = Frontoparietal network; SMN = Sensorimotor network; VAN = Ventral attention network; VN = Visual network.

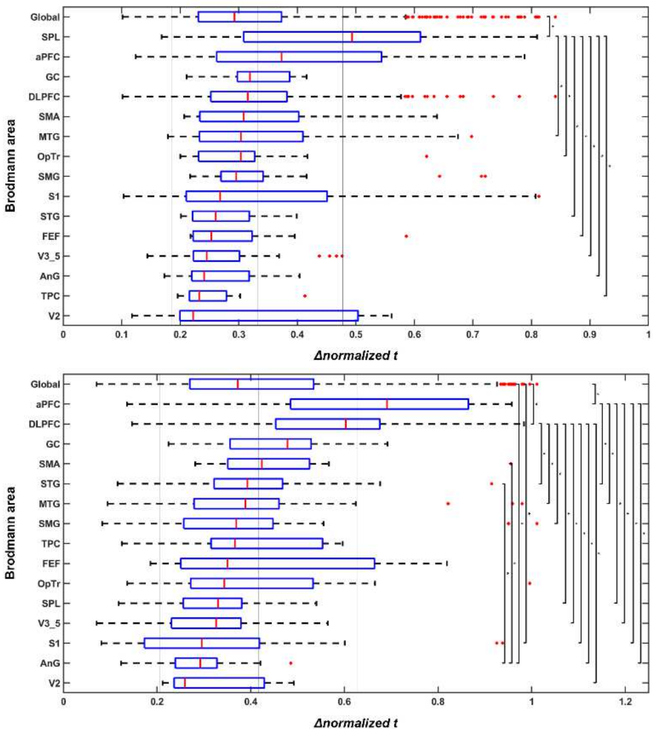


**Fig. S4. Boxplot of the Δnormalized t across the cortex at the induction phase of unconsciousness.** Upper: Range of frequencies < 46 Hz; Lower: Range of (62–150) Hz. * p < 0.05/105. Upper reference line = mean + standard deviation of all channels; Middle reference line = mean of all channels; Lower reference line = mean – standard deviation of all channels. Abbreviations: S1 = Primary somatosensory cortex (BA 1, 2, 3); SMA = Supplementary motor area and premotor cortex (BA 6); SPL = Superior parietal lobule (BA 7); FEF = Frontal eye fields (BA 8); DLPFC = Dorsolateral prefrontal cortex (BA 9, 46); aPFC = Anterior prefrontal cortex (BA 10); V2 = Secondary visual cortex (BA 18); V3–5 = Associative visual cortex (BA 19); MTG = Middle temporal gyrus (BA 21); STG = Superior temporal gyrus (BA 22); TPC = Temporopolar area (BA 38); AnG = Angular gyrus (BA39); SMG = Supramarginal gyrus (BA40); GC = Primary gustatory cortex

(BA 43); OpTr = The opercular part and triangular part of the inferior frontal gyrus (BA 44, 45); Global = all ECoG channels.

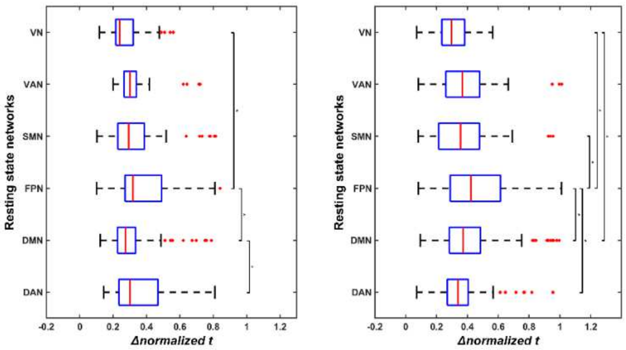


**Fig. S5. Boxplot of the Δnormalized t within resting state networks at the induction phase of unconsciousness.** Left: Range of frequencies < 46 Hz; Right: Range of (62–150) Hz. * p < 0.05/15. Abbreviations: DAN = Dorsal attention network; DMN = Default mode network; FPN = Frontoparietal network; SMN = Sensorimotor network; VAN = Ventral attention network; VN = Visual network.

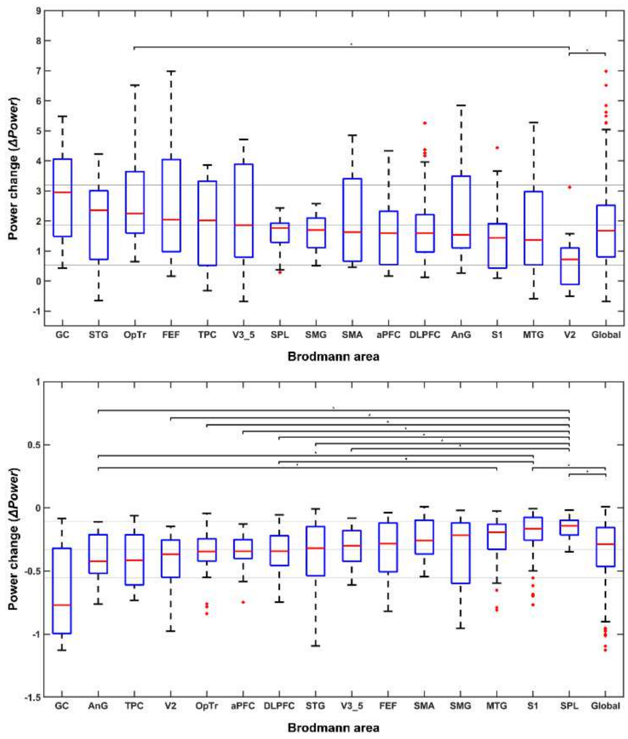


**Fig. S6. Boxplot of the ΔPower across the cortex at the induction phase of unconsciousness.** Upper: Range of frequencies < 46 Hz; Lower: Range of (62–150) Hz. * p < 0.05/105. Upper reference line = mean + standard deviation of all channels; Middle reference line = mean of all channels; Lower reference line = mean – standard deviation of all channels. Abbreviations: S1 = Primary somatosensory cortex (BA 1, 2, 3); SMA = Supplementary motor area and premotor cortex (BA 6); SPL = Superior parietal lobule (BA 7); FEF

= Frontal eye fields (BA 8); DLPFC = Dorsolateral prefrontal cortex (BA 9, 46); aPFC = Anterior prefrontal cortex (BA 10); V2 = Secondary visual cortex (BA 18); V3–5 = Associative visual cortex (BA 19); MTG = Middle temporal gyrus (BA 21); STG = Superior temporal gyrus (BA 22); TPC = Temporopolar area (BA 38); AnG = Angular gyrus (BA39); SMG = Supramarginal gyrus (BA40); GC = Primary gustatory cortex (BA 43); OpTr = The opercular part and triangular part of the inferior frontal gyrus (BA 44, 45); Global = all ECoG channels.

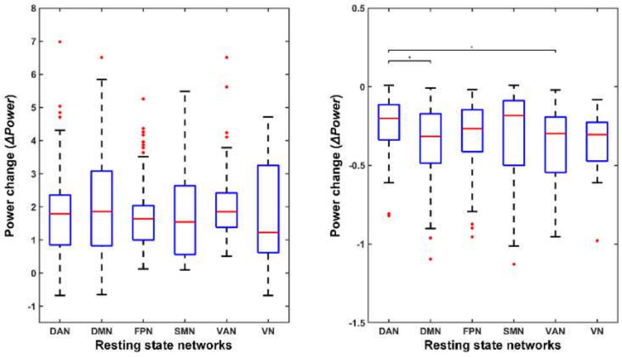


**Fig. S7. Boxplot of the ΔPower within resting state networks at the induction phase of unconsciousness.** Upper: Range of frequencies < 46 Hz; Lower: Range of (62–150) Hz. * $p < 0.05/15$. Abbreviations: DAN = Dorsal attention network; DMN = Default mode network; FPN = Frontoparietal network; SMN = Sensorimotor network; VAN = Ventral attention network; VN = Visual network.

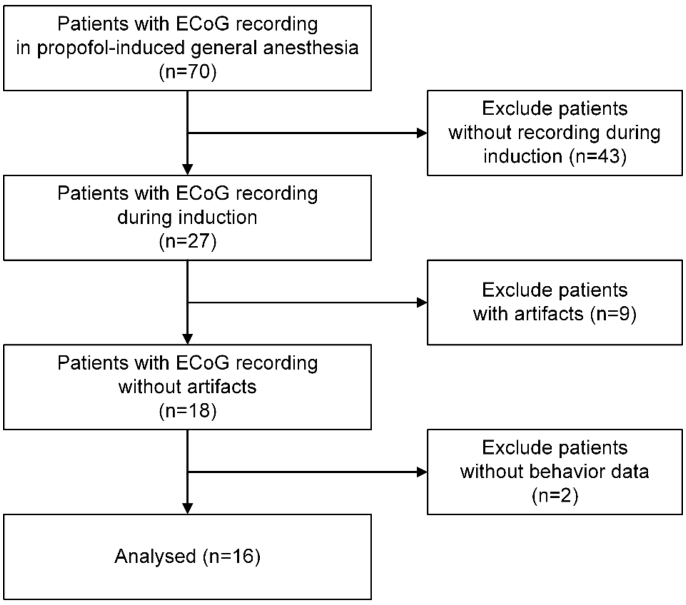


**Fig. S8. Subject flow chart.**

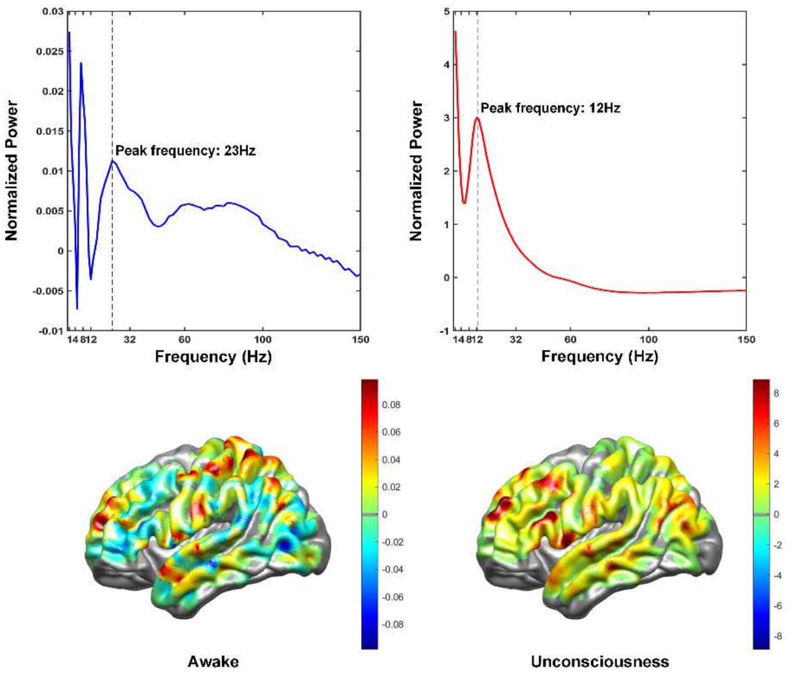


**Fig. S9. The average peak frequency and spatial distribution of power at the peak frequency in the awake and unconsciousness states.**

**Table S1.** Statistical significance of the start point of power change across the cortex at the induction phase of unconsciousness.

| Frequency | Cortical region 1 | Cortical region 2 | Mean difference | P |
|---|---|---|---|---|
| < 46Hz | SPL | Global | SPL < Global | 0.00000 |
| | | S1 | SPL < S1 | 0.00007 |
| | | FEF | SPL < FEF | 0.00002 |
| | | DLPFC | SPL < DLPFC | 0.00005 |
| | | V3–5 | SPL < V3–5 | 0.00000 |
| | | MTG | SPL < MTG | 0.00000 |
| | | STG | SPL < STG | 0.00000 |
| | | TPC | SPL < TPC | 0.00001 |
| | | AnG | SPL < AnG | 0.00000 |
| | | OpTr | SPL < OpTr | 0.00000 |
| | V3–5 | Global | V3–5 > Global | 0.00262 |
| | | DLPFC | V3–5 > DLPFC | 0.00013 |
| | | SMG | V3–5 > SMG | 0.00017 |
| | STG | Global | STG > Global | 0.00034 |
| | | DLPFC | STG > DLPFC | 0.00000 |
| | | SMG | STG > SMG | 0.00001 |
| | | aPFC | STG > aPFC | 0.00031 |
| | AnG | Global | AnG > Global | 0.00098 |

| | | | | |
|---|---|---|---|---|
| | | DLPFC | AnG > DLPFC | 0.00001 |
| | | SMG | AnG > SMG | 0.00003 |
| > 62Hz and < 150Hz | SPL | Global | SPL < Global | 0.00006 |
| | | V3–5 | SPL < V3–5 | 0.00005 |
| | | MTG | SPL < MTG | 0.00008 |
| | | STG | SPL < STG | 0.00004 |
| | | AnG | SPL < AnG | 0.00000 |
| | AnG | Global | AnG > Global | 0.00001 |
| | | DLPFC | AnG > DLPFC | 0.00000 |
| | | aPFC | AnG > aPFC | 0.00010 |
| | | STG | AnG > STG | 0.00004 |
| | | SMG | AnG > SMG | 0.00007 |

**Abbreviations:** S1 = Primary somatosensory cortex (BA 1, 2, 3); SPL = Superior parietal lobule (BA 7); FEF = Frontal eye fields (BA 8); DLPFC = Dorsolateral prefrontal cortex (BA 9, 46); aPFC = Anterior prefrontal cortex (BA 10); V3–5 = Associative visual cortex (V3, V4, V5, BA 19); MTG = Middle temporal gyrus (BA 21); STG = Superior temporal gyrus (BA 22); TPC = Temporopolar area (BA 38); AnG = Angular gyrus (BA39); SMG = Supramarginal gyrus (BA40); OpTr = The opercular part and triangular part of the inferior frontal gyrus (BA 44, 45); Global = all ECoG channels.

**Table S2.** Statistical significance of the Δnormalized t across the cortex at the induction phase of unconsciousness.

| Frequency | Cortical region 1 | Cortical region 2 | Mean difference | P |
|---|---|---|---|---|
| < 46Hz | SPL | Global | SPL > Global | 0.00000 |
| | | FEF | SPL > FEF | 0.00005 |
| | | V3–5 | SPL > V3–5 | 0.00000 |
| | | MTG | SPL > MTG | 0.00003 |
| | | STG | SPL > STG | 0.00000 |
| | | TPC | SPL > TPC | 0.00002 |
| | | AnG | SPL > AnG | 0.00000 |
| | | OpTr | SPL > OpTr | 0.00030 |
| > 62Hz and < 150Hz | DLPFC | Global | DLPFC > Global | 0.00000 |
| | | S1 | DLPFC > S1 | 0.00000 |
| | | SPL | DLPFC > SPL | 0.00000 |
| | | V2 | DLPFC > V2 | 0.00027 |
| | | V3–5 | DLPFC > V3–5 | 0.00000 |
| | | MTG | DLPFC > MTG | 0.00000 |
| | | STG | DLPFC > STG | 0.00000 |
| | | AnG | DLPFC > AnG | 0.00000 |
| | | SMG | DLPFC > SMG | 0.00006 |
| | aPFC | Global | aPFC > Global | 0.00010 |
| | | S1 | aPFC > S1 | 0.00004 |
| | | SPL | aPFC > SPL | 0.00002 |
| | | V3–5 | aPFC > V3–5 | 0.00004 |

| | | | | |
|---|---|---|---|---|
| | | MTG | aPFC > MTG | 0.00028 |
| | | STG | aPFC > STG | 0.00021 |
| | | AnG | aPFC > AnG | 0.00000 |
| | AnG | Global | AnG < Global | 0.00007 |
| | | SMA | AnG < SMA | 0.00006 |
| | | STG | AnG < STG | 0.00001 |
| | S1 | Global | S1 < Global | 0.00211 |

**Abbreviations:** S1 = Primary somatosensory cortex (BA 1, 2, 3); SMA = Supplementary motor area and premotor cortex (BA 6); SPL = Superior parietal lobule (BA 7); FEF = Frontal eye fields (BA 8); DLPFC = Dorsolateral prefrontal cortex (BA 9, 46); aPFC = Anterior prefrontal cortex (BA 10); V2 = Secondary visual cortex (BA 18); V3–5 = Associative visual cortex (BA 19); MTG = Middle temporal gyrus (BA 21); STG = Superior temporal gyrus (BA 22); TPC = Temporopolar area (BA 38); AnG = Angular gyrus (BA39); SMG = Supramarginal gyrus (BA40); OpTr = The opercular part and triangular part of the inferior frontal gyrus (BA 44, 45).

**Table S3.** Statistical significance of the ΔPower across the cortex at the induction phase of unconsciousness.

| Frequency | Cortical region 1 | Cortical region 2 | Mean difference | P |
|---|---|---|---|---|
| < 46Hz | V2 | Global | V2 < Global | 0.00248 |
| | | OpTr | V2 < OpTr | 0.00023 |
| > 62Hz and < 150Hz | S1 | Global | S1 > Global | 0.00053 |
| | | DLPFC | S1 > DLPFC | 0.00004 |
| | | AnG | S1 > AnG | 0.00013 |
| | SPL | Global | SPL > Global | 0.00000 |
| | | DLPFC | SPL > DLPFC | 0.00000 |
| | | aPFC | SPL > aPFC | 0.00000 |
| | | V2 | SPL > V2 | 0.00008 |
| | | V3–5 | SPL > V3–5 | 0.00007 |
| | | STG | SPL > STG | 0.00007 |
| | | AnG | SPL > AnG | 0.00000 |
| | | OpTr | SPL > OpTr | 0.00001 |
| | AnG | MTG | AnG < MTG | 0.00044 |

**Abbreviations:** S1 = Primary somatosensory cortex (BA 1, 2, 3); SPL = Superior parietal lobule (BA 7); DLPFC = Dorsolateral prefrontal cortex (BA 9, 46); aPFC = Anterior prefrontal cortex (BA 10); V2 = Secondary visual cortex (BA 18); V3–5 = Associative visual cortex (V3, V4, V5, BA 19); MTG = Middle temporal gyrus (BA 21); STG = Superior temporal gyrus (BA 22); AnG = Angular gyrus (BA39); OpTr = The opercular part and triangular part of the inferior frontal gyrus (BA 44, 45); Global = all ECoG channels.

**Table S4.** Demographics and clinical characteristics of patients.

| Subject | Age | Sex | Handness | Number of Electrodes | Electrode location | Diagnosis | Time to loss of consciousness (sec) |
|---|---|---|---|---|---|---|---|
| Sub 1 | 58 | F | R | 14 | R | Temporal lobe epilepsy | 247 |
| Sub 2 | 64 | M | Mixed | 5 | R | Temporal lobe epilepsy | 277 |
| Sub 3 | 19 | M | R | 6 | L | Temporal lobe epilepsy | 120 |

| | | | | | | | |
|---|---|---|---|---|---|---|---|
| Sub 4 | 30 | M | R | 8 | R | Temporal lobe epilepsy | 179 |
| Sub 5 | 24 | M | R | 24 | L | Parietal lobe epilepsy | 176 |
| Sub 6 | 40 | F | R | 35 | R | Frontal lobe epilepsy | 199 |
| Sub 7 | 36 | F | R | 43 | L | Frontal lobe epilepsy | 233 |
| Sub 8 | 41 | M | R | 19 | R | Temporal lobe epilepsy | 98 |
| Sub 9 | 29 | M | R | 28 | L | Frontal lobe epilepsy | 346 |
| Sub 10 | 32 | F | R | 14 | L | Temporal lobe epilepsy | 158 |
| Sub 11 | 22 | F | R | 42 | R | Temporal lobe epilepsy | 125 |
| Sub 12 | 23 | F | R | 39 | R | Temporal lobe epilepsy | 151 |
| Sub 13 | 24 | M | R | 37 | L | Parietal lobe epilepsy | 240 |
| Sub 14 | 35 | F | R | 48, 53* | L | Parietal lobe epilepsy | 158, 204 |
| Sub 15 | 25 | M | R | 26 | R | Frontal lobe epilepsy | 247 |
| Sub 16 | 27 | F | R | 23 | L | Temporal lobe epilepsy | 159 |
| **Mean** | **33.1** | | | | | | **195.1** |

**Abbreviations:** M = Male; F = Female; R = Right; L = Left; * The patient underwent additional electrode insertion to different locations.

**Table S5.** Distribution of electrodes.

| Brodmann area | Abbreviation | Name of the Brodmann area | Number of Subjects | Number of Electrodes |
|---|---|---|---|---|
| **BA 1, 2, 3** | S1 | Primary somatosensory cortex | 9 | 37 |
| **BA 6** | SMA | Supplementary motor area and premotor cortex | 7 | 14 |
| **BA 7** | SPL | Superior parietal lobule | 5 | 33 |
| **BA 8** | FEF | Frontal eye fields | 5 | 17 |
| **BA 9, 46** | DLPFC | Dorsolateral prefrontal cortex | 7 | 75 |
| **BA 10** | aPFC | Anterior prefrontal cortex | 5 | 21 |
| **BA 18** | V2 | Secondary visual cortex | 3 | 11 |
| **BA 19** | V3–5 | Associative visual cortex | 6 | 27 |
| **BA 21** | MTG | Middle temporal gyrus | 10 | 47 |
| **BA 22** | STG | Superior temporal gyrus | 8 | 45 |
| **BA 38** | TPC | Temporopolar area | 5 | 13 |
| **BA 39** | AnG | Angular gyrus | 5 | 36 |
| **BA 40** | SMG | Supramarginal gyrus | 7 | 31 |
| **BA 43** | GC | Primary gustatory cortex | 4 | 11 |
| **BA 44, 45** | OpTr | The opercular part and triangular part of the inferior frontal gyrus | 7 | 22 |
| **All** | | | **17** | **440** |
